# Six novel nsSNPs affect *RUNX1* gene may leading to Acute Myeloid Leukemia (AML) using Bioinformatics approach

**DOI:** 10.1101/2023.06.26.546458

**Authors:** Dalia Omer, Tebyan Ameer, Tarig Alsheikh, Ahmed NjmEldin, Zeinab Abdalmonem, Hadil Suliman, Abubaker Agbash, Howaida Hamad, Baha Aldeen Alshareif, Mohammed A. Hassan

## Abstract

**Background:** *RUNX1* is one of the most frequently mutated genes in human AMLs, most of *RUNX1* mutations in acute myeloid leukemia (AML) are missense or deletion-truncation and behave as loss-of-function mutations. The molecular consequences of cancer associated mutations in Acute myeloid leukemia (AML) linked factors are not very well understood. Here, we recognize possible pathogenic SNPs in the *RUNX1* gene as Functional differences caused by SNPs might have harmful effects on protein structure and function using various computational tools.

**Methodology:** Data gained from NCBI database and various tools used to study nsSNPs which they are: SIFT, Polyphen-2, Provean, SNAP2, I-Mutant, Project Hope, Raptor X, PolymiRTS and Gene MANIA.

**Result:** Our study reveals six novel SNPs observed to be the most damaging SNPs that affect structure and function of *RUNX1* gene using various bioinformatics tools.

**Conclusion:** This study revealed 7 damaging SNPs, 6 novel nsSNP out of them in the *RUNX1* gene that leads to AML, by using different bioinformatics tools. Also, 23 functional classes were predicted in 8 SNPs in the 3’UTR, among them, 6 alleles disrupted a conserved miRNA site and 16 derived alleles created a new site of miRNA. This might result in the de regulation of the gene function. Hopefully, these results will help in genetic studying and diagnosis of AML improvement.

## INTRODUCTION

Leukemia is thought to develop by a variety of biological pathways, including a block in the differentiation or maturation of hematopoietic progenitors, an increase in the lifespan of myeloid or lymphoid cells, and uncontrolled proliferation of hematopoietic cell (1).

The two groups of mutations that are thought to induce leukemia are class I mutations, which are thought to cause unregulated proliferation or impede apoptosis of the cells, and class II mutations, which are thought to cause block in development or maturation of the cells. Class I mutations include those causing abnormal tyrosine kinase activity such as BCR-ABL in chronic myelogenous leukemia and constitutive expression of antiapoptotic signalling molecules such as bcl-2 and Myc in B cell lymphomas. Since loss of *RUNX1* function has been supposed to cause block in the differentiation of hematopoietic cells, *RUNX1* mutations have been recognized as class II mutations (2).

The development of acute myeloid leukemia (AML) results from the recurrent acquisition of gene mutations in hematopoietic progenitor cells, these modifications impair healthy hematopoietic growth and differentiation, causing a differentiation block that accumulates aberrant and immature myeloid cells (blasts) in the bone marrow, peripheral blood and occasionally extramedullary sites(3).

*RUNX1* is one of the most frequently mutated genes in human AMLs, most of *RUNX1* mutations in acute myeloid leukemia (AML) are missense or deletion-truncation and behave as loss-of-function mutations (4).

*RUNX1* mutations are discovered in roughly 5.6–17.9% of AML cases, 3% of individuals with pediatric AML, and about 27.7% of secondary AML that has undergone MDS transformation. Besides being associated with older age and male gender, the frequency of *RUNX1* mutation was reported to vary in different risk levels of patients and French-American-British (FAB) subtypes (5).

*RUNX1* gene located on chromosome 21, also known as acute myeloid leukemia 1 protein (AML1), core-binding factor subunit alpha-2 (CBFA2) or polyoma virus enhancer-binding protein 2αB (PEBP2αB) (6), is a transcription factor, controlling the differentiation of hematopoietic stem cells into mature blood cells. Also, it’s one of the most frequently likened genes in a variety of haematological malignancies. *RUNX1* dysfunction results from either gene translocations, point mutations, or aberrant expression and is a major predisposing risk factor for developing haematological malignancies (7).

*RUNX1* mutations in leukemias were initially reported as chimeric genes formed by chromosomal translocations including t(8;21), t(3;21), and t(16;21)(8). Afterwards, *RUNX1* mutations were also found in the myeloid malignancies with normal karyotype, including FAB classification M0 AMLs and myelodysplastic syndromes (MDS). In the FAB M0 AMLs, *RUNX1* mutations were detected in more than 40 % of cases, and *RUNX1* point mutations are also detected in patients with myeloproliferative neoplasms (MPNs) transformed into AMLs. Presence of *RUNX1* mutations independently predicted worse prognosis and resistance to chemotherapy in both younger and older patients with CN-AML treated with intensive induction chemotherapy and not receiving alloSCT. Thus, patients harbouring *RUNX1* mutations warrant strong consideration of up-front novel therapies and/or early alloSCT (9, 10).

The molecular consequences of cancer associated mutations in Acute myeloid leukemia (AML) linked factors are not very well understood. Here, we interrogated the database for missense mutations associated with the *RUNX1* protein, that is frequently miss-regulated in AML (11).

Science will advance more quickly and effectively if bioinformatics resources and computational “In-Silico” experiments are combined. So, in the current study, we sought to ascertain the impact of *RUNX1* gene polymorphisms on the protein structure/function of the gene, which may play a significant role in disease susceptibility.

## MATERIALS AND METHODS

Data on *RUNX1* gene was obtained from the national center for biological information (NCBI) website (https://www.ncbi.nlm.nih.gov/) and the SNPs (single nucleotide polymorphisms) information was retrieved from NCBI SNPs database dbSNP (https://www.ncbi.nlm.nih.gov/snp/). The gene ID and sequence was obtained from Uiprot (https://www.uniprot.org/). Analysis of the SNPs was done according to figure 1.

**Figure 1.**
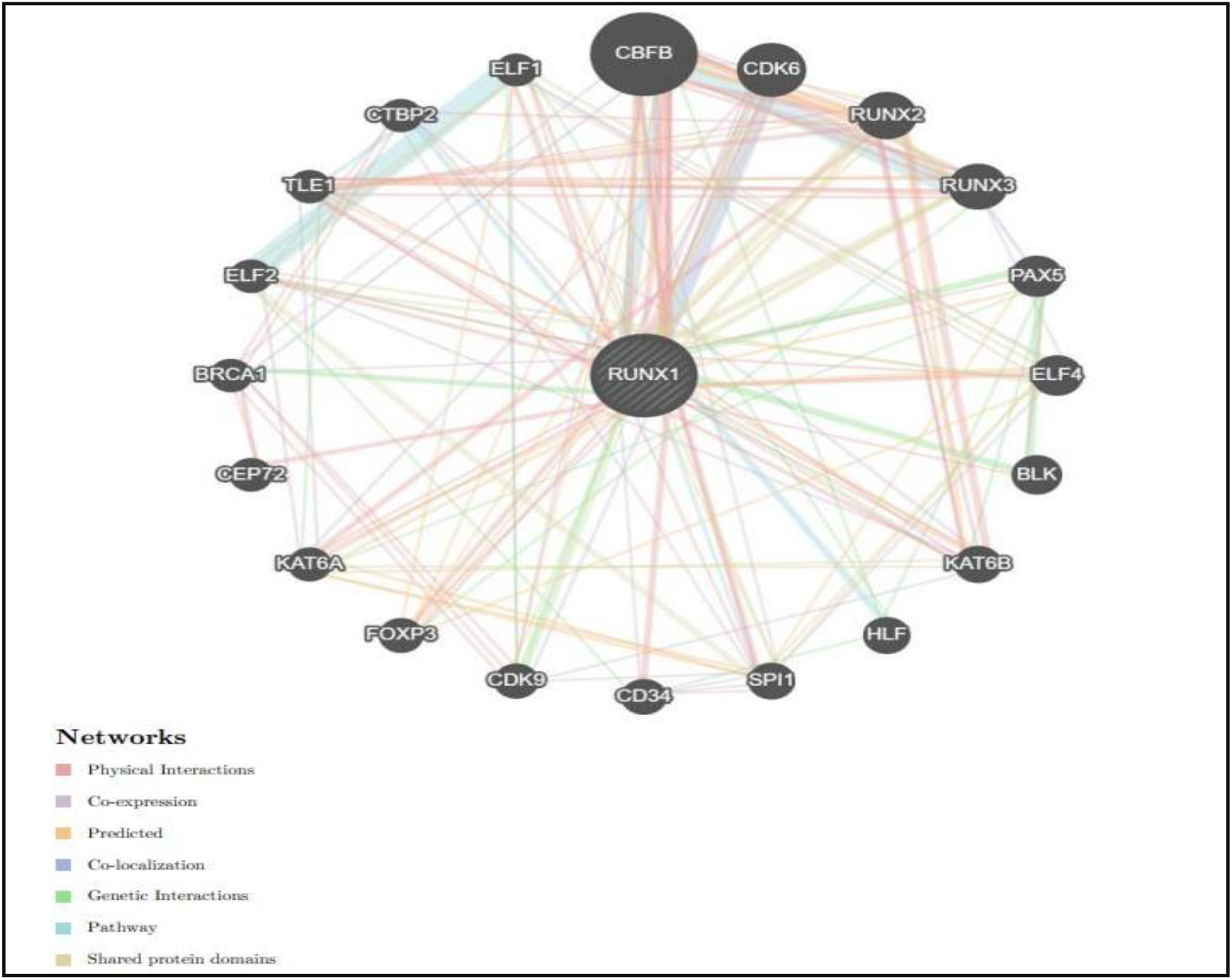
*RUNX1* Gene Interactions and network predicted by GeneMania.

**Figure 2:**
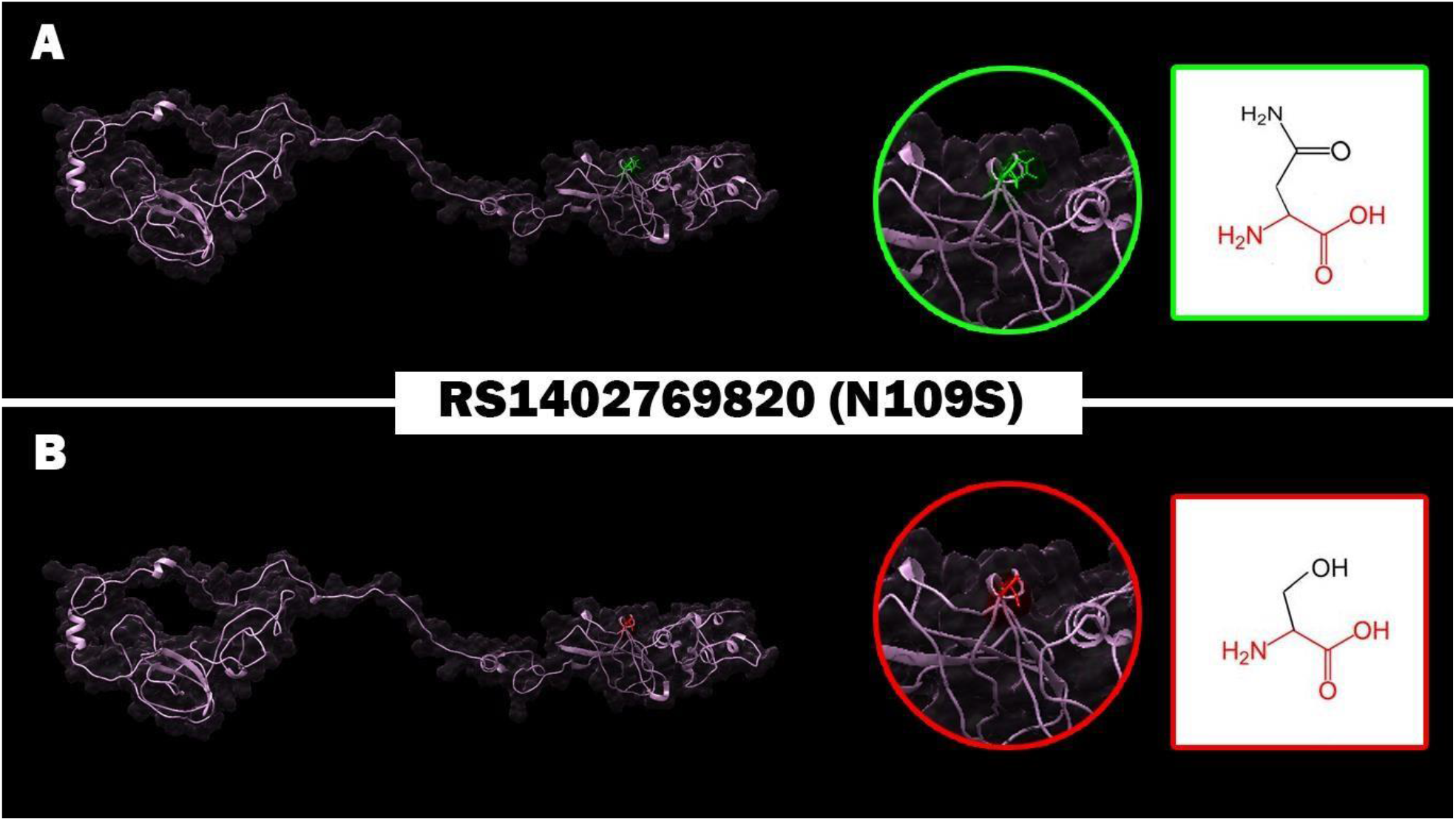
Effect of N109S (rs1402769820) SNP on protein structure in which Asparagine (Green residue) mutated into Serine (Red residue) at position 109.

**Figure 3:**
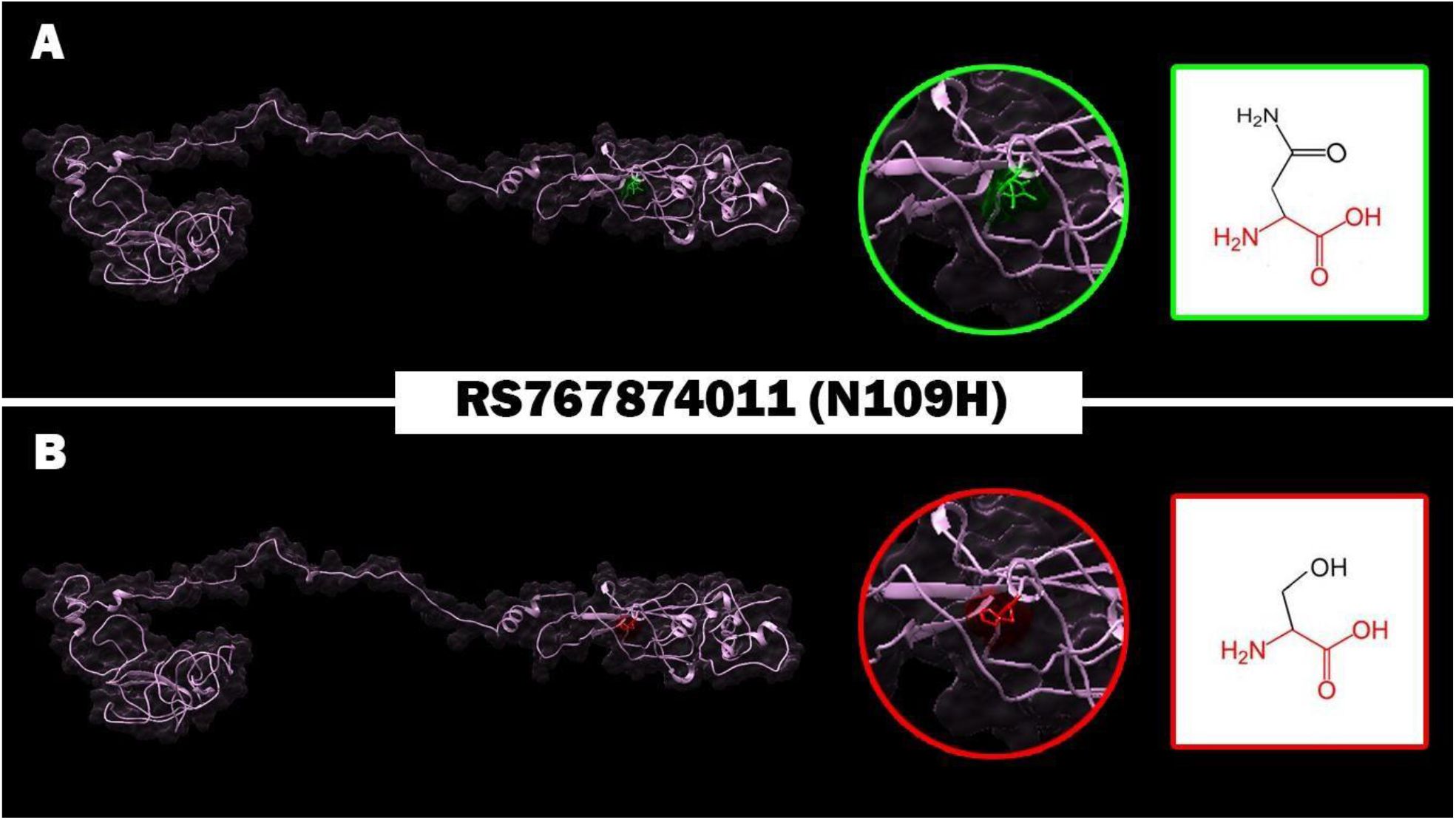
Effect of N109S (rs767874011) SNP on protein structure in which Asparagine (Green residue) mutated into Histidine (Red residue) at position 109.

**Figure 4:**
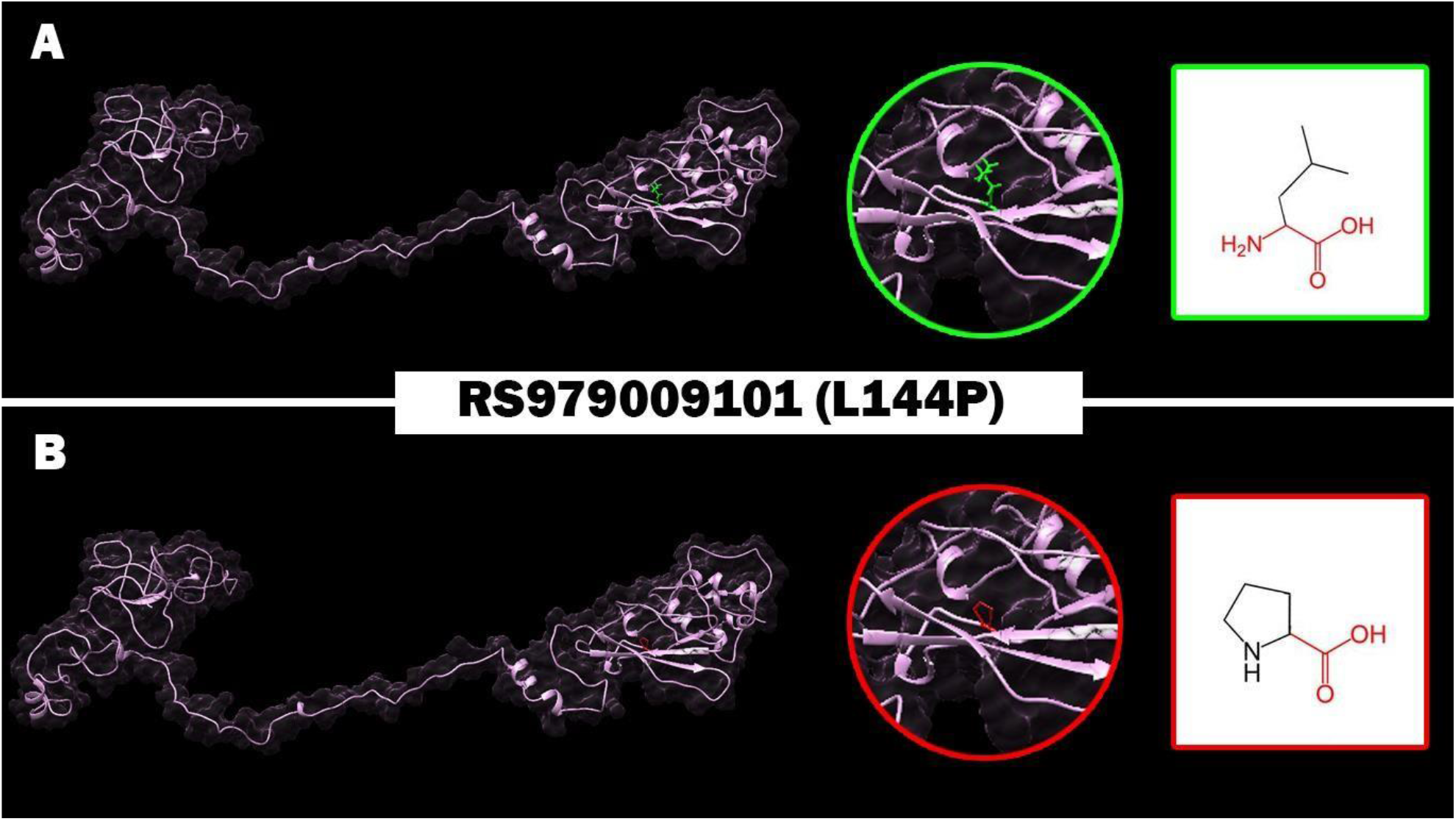
Effect of L144P (rs979009101) SNP on protein structure in which Leucine (Green residue) mutated into Proline (Red residue) at position 144.

**Figure 5:**
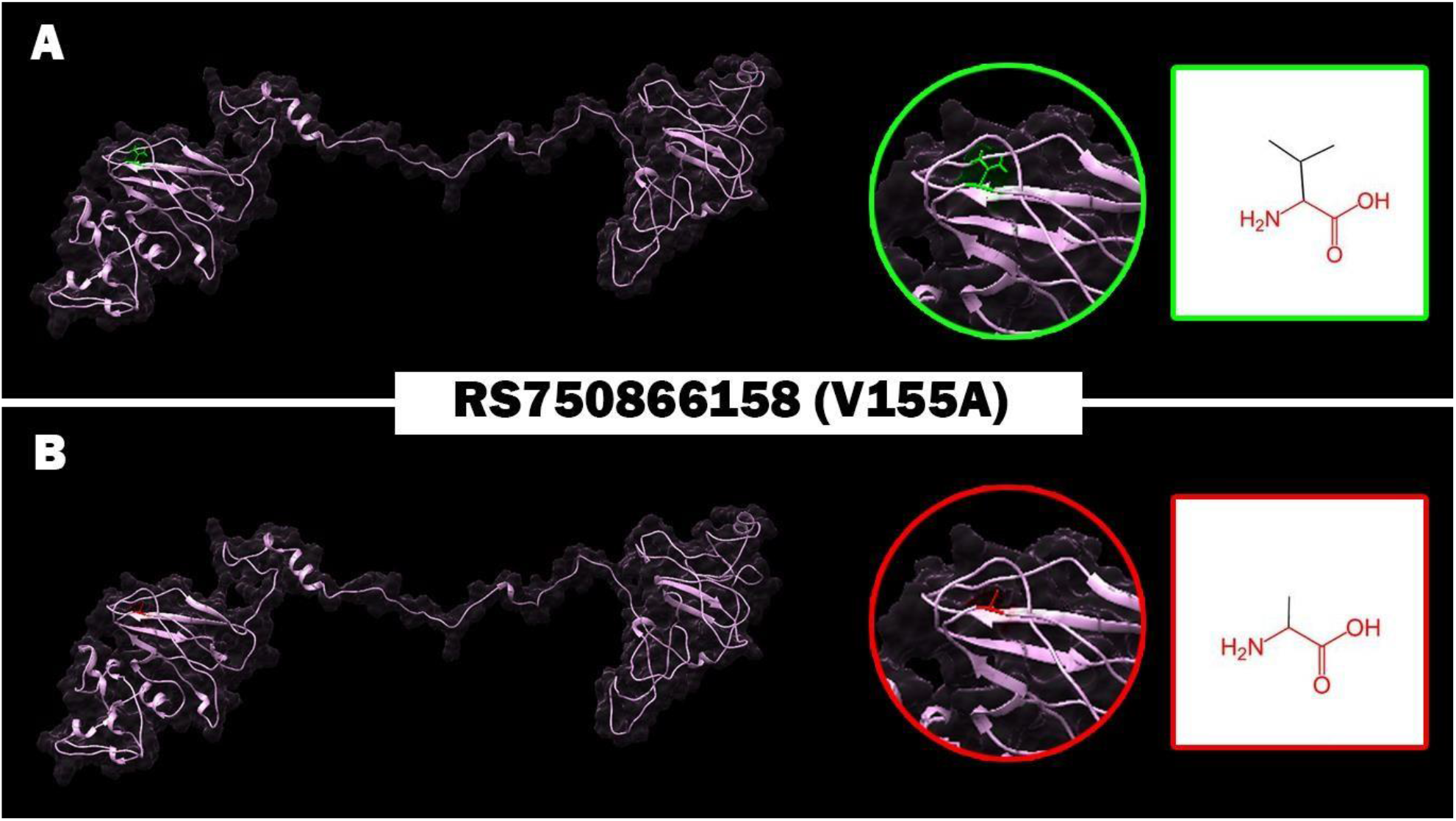
Effect of V155A (rs750866158) SNP on protein structure in which Valine (Green residue) mutated into Alanine (Red residue) at position 155.

**Figure 6:**
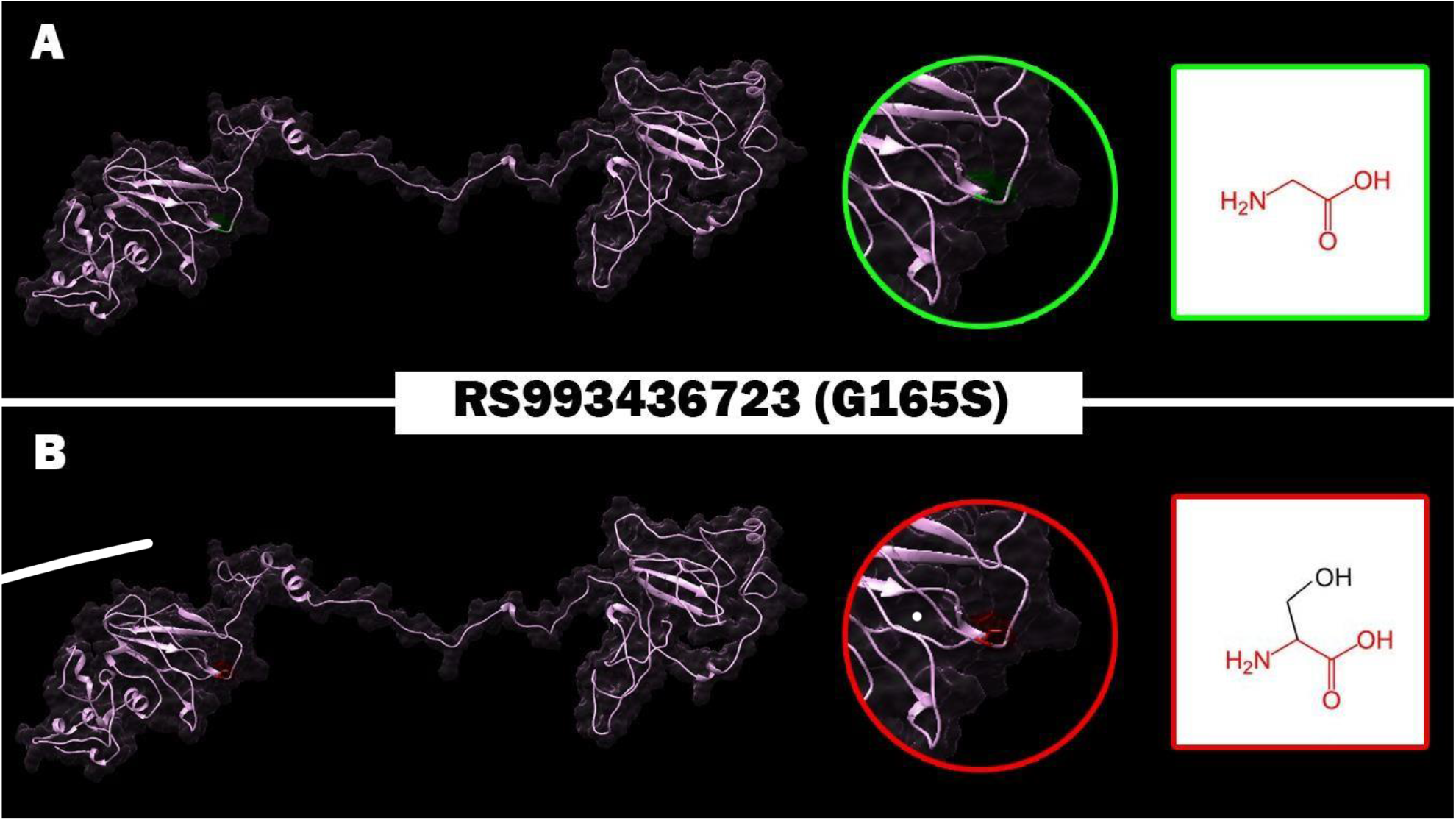
Effect of G165S (rs993436723) SNP on protein structure in which Glycine (Green residue) mutated into Serine (Red residue) at position 165.

**Figure 7:**
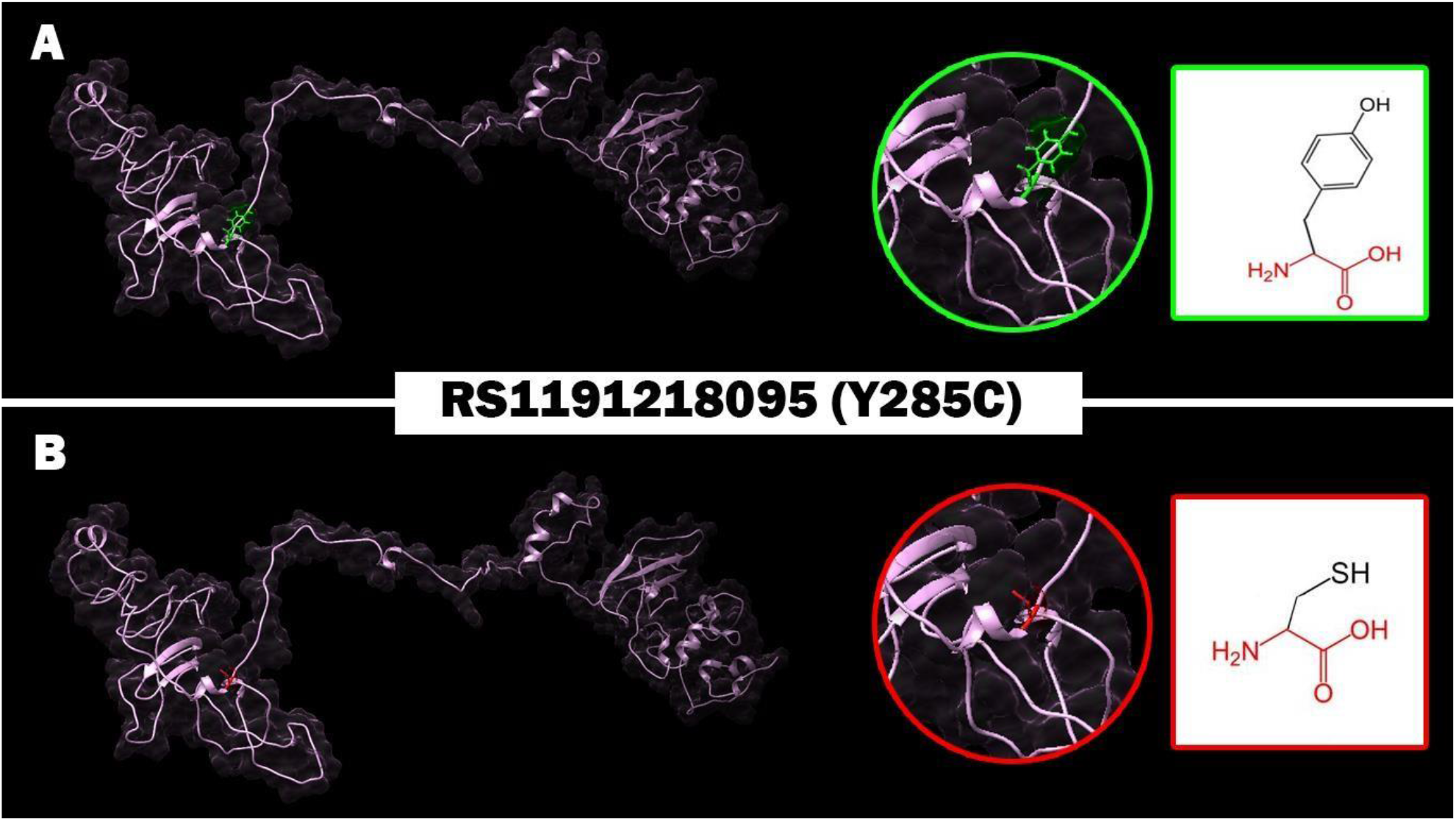
Effect of Y285C (rs1191218095) SNP on protein structure in which Tyrosine (Green residue) mutated into Cysteine (Red residue) at position 285.

**Figure 8:**
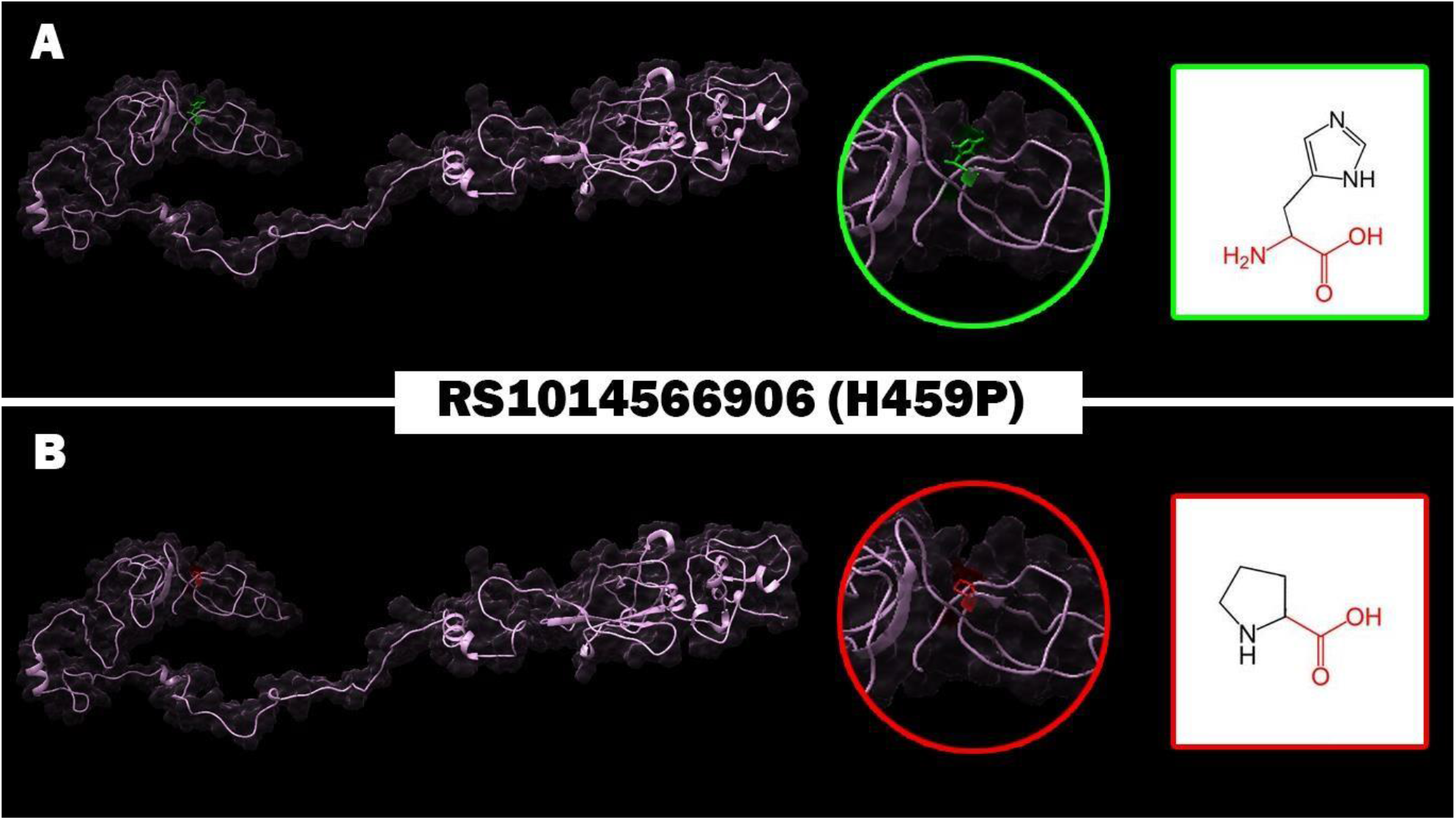
Effect of H459P (rs1191218095) SNP on protein structure in which Histidine (Green residue) mutated into Proline (Red residue) at position 459.

### SIFT

(Sorting Intolerant from Tolerant) is a sequence homology-based tool that sorts intolerant from tolerant amino acid substitutions and predicts whether an amino acid substitution in a protein will have a phenotypic outcome, considering the position at which the mutation occurred and the type of amino acid change. When a protein sequence submitted, SIFT chooses related proteins and obtains an alignment of these proteins with the query. Based on the change on the type of amino acids appearing at each position in the alignment, SIFT calculates the probability that amino acid at a position is tolerated conditional on the most frequent amino acid being tolerated. If this normalized value is less than a cut-off, the substitution is predicted to be deleterious. SIFT scores 0.05 are considered tolerant. Available at (http://sift.bii.a-star.edu.sg/) (12).

### PolyPhen-2

Polymorphism Phenotyping v.2 available at (http://genetics.bwh.harvard.edu/pph2/) is a tool that predicts the possible effect of an amino acid substitution on the structure and function of a human protein by using simple physical and comparative considerations. The submission of the sequence allows querying for a single individual amino acid substitution or a coding, non-synonymous SNP annotated in the SNP database. It calculates position-specific independent count (PSIC) scores for each of the two variants and computes the difference of the PSIC scores of the two variants. The higher a PSIC score difference, the higher the functional effect of a particular amino acid substitution is likely to have. PolyPhen scores were designated as probably damaging (0.95–1), possibly damaging (0.7–0.95), and benign (0.00– 0.31) (13).

### Provean

Protein Variation Effect Analyzer software available at (http://provean.jcvi.org/index.php) it predicts whether an amino acid substitution has an impact on the biological function of the protein. Provean is useful in filtrating sequence variants to identify non-synonymous variants that are predicted to be functionally important (14).

### SNAP2

Available at (https://rostlab.org/owiki/index.php/Snap2), SNAP2 is a trained classifier that is based on a machine learning device called “neural network”. It distinguishes between effect and neutral variants/non-synonymous SNPs by taking a variety of sequence and variant features in considerations. The most important input signal for the prediction is the evolutionary information taken from an automatically generated multiple sequence alignment. Also, structural features such as predicted secondary structure and solvent accessibility are considered. If available also annotation (i.e. known functional residues, pattern, regions) of the sequence or close homologs are pulled in. SNAP2 has persistent two states accuracy (effect/neutral) of 82 % (15).

### SNP & GO

It is a server for the prediction of single point protein mutations likely to be involved in the causing of diseases in humans. (https://snps-and-go.biocomp.unibo.it/snps-and-go/) (16).

### PHD-SNP

Prediction of human Deleterious Single Nucleotide Polymorphisms is available at (http://snps.biofold.org/phd-snp/phd-snp.html). The working principle is Support Vector Machines (SVMs) based method trained to predict disease-associated nsSNPs using sequence information. The related mutation is predicted as disease-related (Disease) or as neutral polymorphism (Neutral) (16).

### I-MUTANT

I-mutant is a suit of support vector machine-based predictors which integrated into a unique web server. It offers the opportunity to predict automatically protein stability changes upon single site mutations starting from protein sequence alone or protein structure if available. Also, it gives the possibility to predict human deleterious SNPs from the protein sequence alone. It is available at (http://gpcr.biocomp.unibo.it/~emidio/I-Mutant3.0/old/IntroIMutant3.0_help.html) (17).

### Project Hope

Online software is available at: (http://www.cmbi.ru.nl/hope/method/). It is a web service where the user can submit a sequence and mutation. The software collects structural information from a series of sources, including calculations on the 3D protein structure, sequence annotations in UniProt and prediction from other software. It combines this information to give analysis for the effect of a certain mutations on the protein structure. HOPE will show the effect of that mutation in such a way that even those without a bioinformatics background can understand it. It allows the user to submit a protein sequence (can be FASTA or not) or an accession code of the protein of interest. In the next step, the user can indicate the mutated residue with a simple mouse click. In the final step, the user can simply click on one of the other 19 amino acid types that will become the mutant residue, and then full report well is generated (18).

### UCSF Chimera (University of California at San Francisco)

UCSF Chimera (https://www.cgl.ucsf.edu/chimera/) is a highly extensible program for interactive visualization and analysis of molecular structures and related data, including density maps, supramolecular assemblies, sequence alignments, docking results, trajectories, and conformational ensembles. High-quality images and animations can be generated. Chimera includes complete documentation and several tutorials. Chimera is developed by the Resource for Biocomputing, Visualization, and Informatics (RBVI), supported by the National Institutes of Health (P41-GM103311) (19).

### PolymiRTS

PolymiRTS is a software used to predict 3UTR (un-translated region) polymorphism in microRNAs and their target sites available at (http://compbio.uthsc.edu/miRSNP/). It is a database of naturally occurring DNA variations in microRNAs (miRNA) seed region and miRNA target sites. MicroRNAs pair to the transcript of protein coding genes and cause translational repression or mRNA destabilization. SNPs in microRNA and their target sites may affect miRNA-mRNA interaction, causing an effect on miRNA-mediated gene repression, PolymiRTS database was created by scanning 3UTRs of mRNAs in human and mouse for SNPs in miRNA target sites. Then, the effect of polymorphism on gene expression and phenotypes are identified and then linked in the database. The PolymiRTS data base also includes polymorphism in target sites that have been supported by a variety of experimental methods and polymorphism in miRNA seed regions (20).

### GeneMANIA

It is gene interaction software that finds other genes which is related to a set of input genes using a very large set of functional association data. Association data include protein and genetic interactions, pathways, co-expression, co-localization and protein domain similarity. GeneMANIA also used to find new members of a pathway or complex, find additional genes you may have missed in your screen or find new genes with a specific function, such as protein kinases. available at (https://genemania.org/) (21).

## RESULTS

### Data Retrieval

Data of human *RUNX1* gene were collected from the National Center for Biological Information (NCBI) website (22). The SNP information (SNP ID) of the *RUNX1* gene was retrieved from the NCBI dbSNP (http://www.ncbi.nlm.nih.gov/snp/) and the protein ID and its sequence was collected from Swiss-Prot databases with the accession number: (Q01196). (http://expasy.org/) (23).

### Functional analysis of nsSNP effect on *RUNX1* using SIFT, Polyphen2, PROVEAN and SNAP2 softwares

**Table 1.**
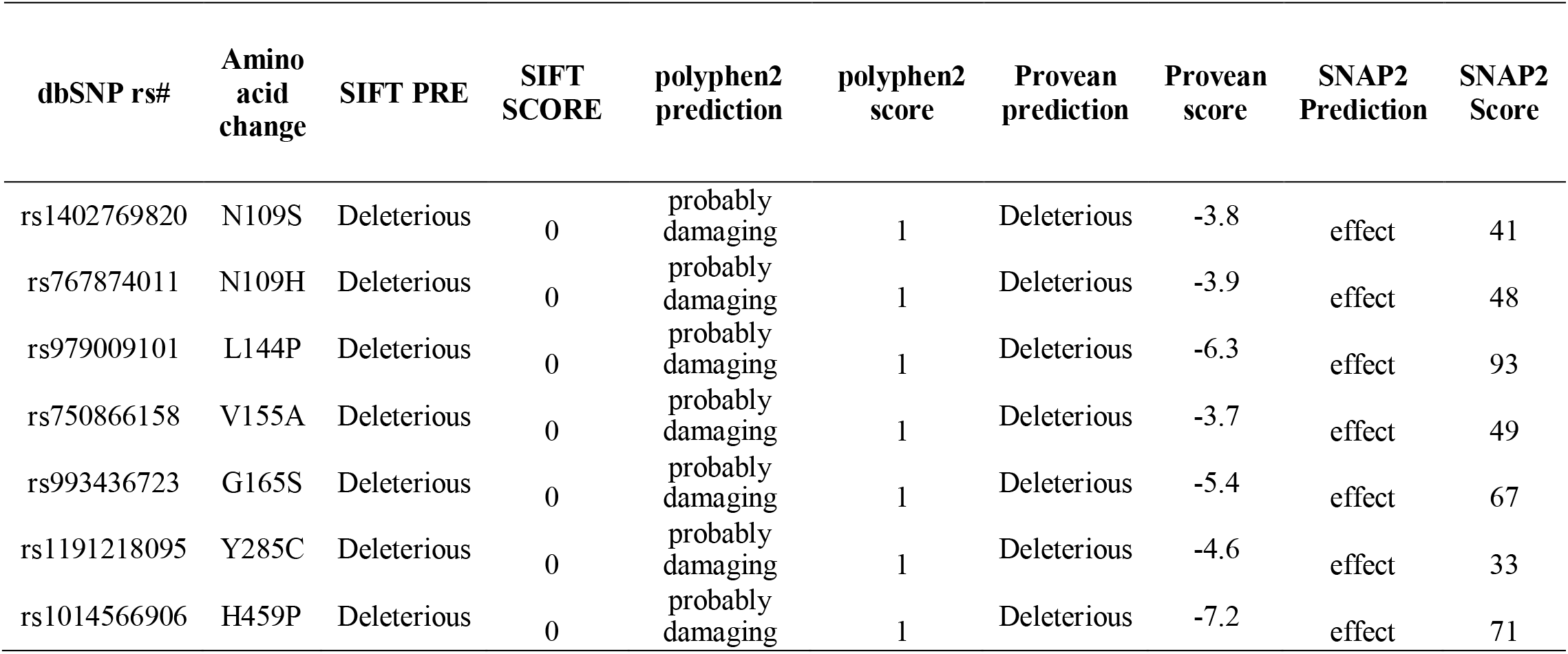
Damaging or Deleterious or effect nsSNPs associated variations predicted by SIFT, Polyphen, PROVEAN, SNAP2 softwares

| dbSNP rs# | Amino acid change | SIFT PRE | SIFT SCORE | polyphen2 prediction | polyphen2 score | Provean prediction | Provean score | SNAP2 Prediction | SNAP2 Score |
| --- | --- | --- | --- | --- | --- | --- | --- | --- | --- |
| rs1402769820 | N109S | Deleterious | 0 | probably damaging | 1 | Deleterious | -3.8 | effect | 41 |
| rs767874011 | N109H | Deleterious | 0 | probably damaging | 1 | Deleterious | -3.9 | effect | 48 |
| rs979009101 | L144P | Deleterious | 0 | probably damaging | 1 | Deleterious | -6.3 | effect | 93 |
| rs750866158 | V155A | Deleterious | 0 | probably damaging | 1 | Deleterious | -3.7 | effect | 49 |
| rs993436723 | G165S | Deleterious | 0 | probably damaging | 1 | Deleterious | -5.4 | effect | 67 |
| rs1191218095 | Y285C | Deleterious | 0 | probably damaging | 1 | Deleterious | -4.6 | effect | 33 |
| rs1014566906 | H459P | Deleterious | 0 | probably damaging | 1 | Deleterious | -7.2 | effect | 71 |

### Disease effect & stability analysis of nsSNP on *RUNX1* using P-Mut & I-Mutant softwares respectively

**Table 2:**
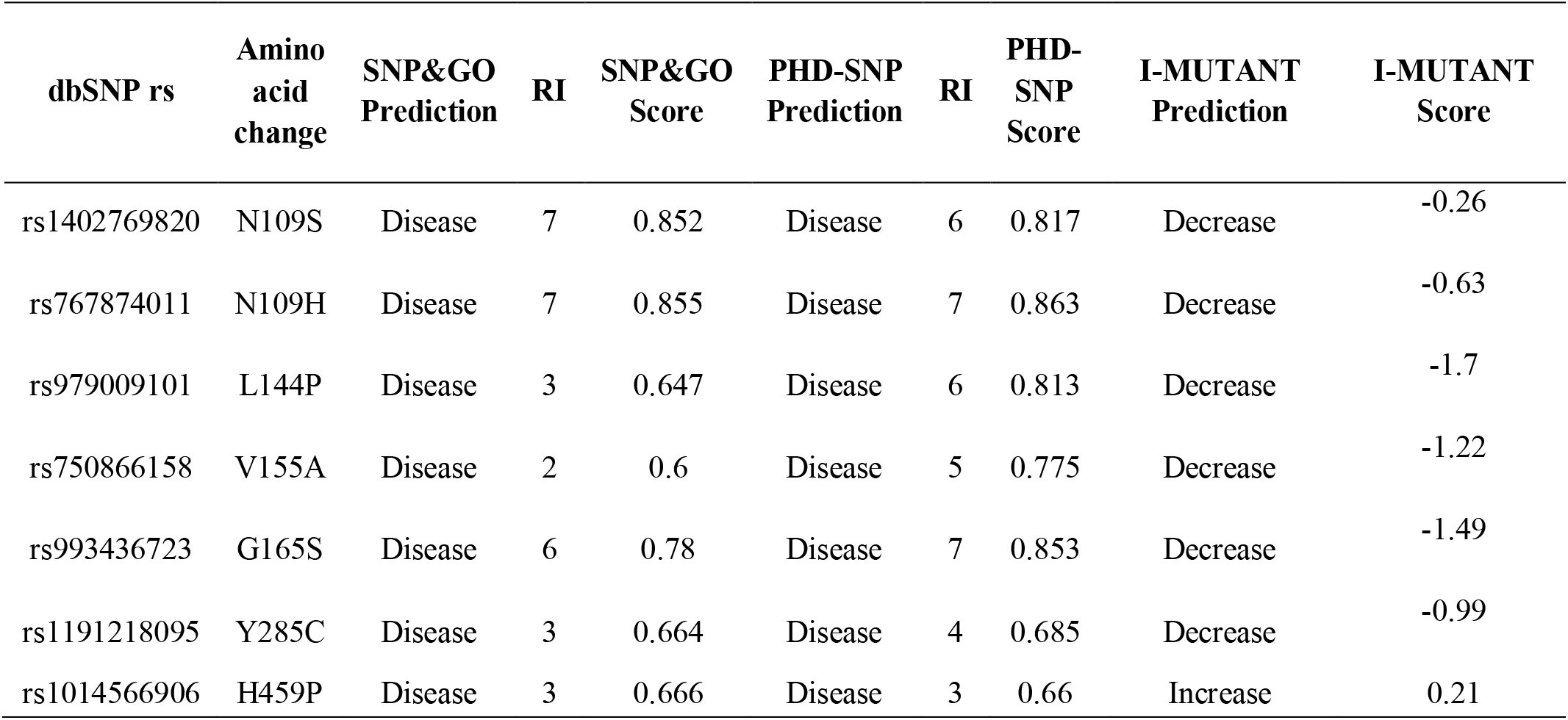
Disease effect & stability analysis of nsSNPs associated variations predicted by SNP & GO, PHD-SNP softwares

| dbSNP rs | Amino acid change | SNP&GO Prediction | RI | SNP&GO Score | PHD-SNP Prediction | RI | PHD-SNP Score | I-MUTANT Prediction | I-MUTANT Score |
| --- | --- | --- | --- | --- | --- | --- | --- | --- | --- |
| rs1402769820 | N109S | Disease | 7 | 0.852 | Disease | 6 | 0.817 | Decrease | -0.26 |
| rs767874011 | N109H | Disease | 7 | 0.855 | Disease | 7 | 0.863 | Decrease | -0.63 |
| rs979009101 | L144P | Disease | 3 | 0.647 | Disease | 6 | 0.813 | Decrease | -1.7 |
| rs750866158 | V155A | Disease | 2 | 0.6 | Disease | 5 | 0.775 | Decrease | -1.22 |
| rs993436723 | G165S | Disease | 6 | 0.78 | Disease | 7 | 0.853 | Decrease | -1.49 |
| rs1191218095 | Y285C | Disease | 3 | 0.664 | Disease | 4 | 0.685 | Decrease | -0.99 |
| rs1014566906 | H459P | Disease | 3 | 0.666 | Disease | 3 | 0.66 | Increase | 0.21 |

### Modelling of amino acid substitution effects on protein structure using Project Hope and Chimera Softwares

### Interactions of *RUNX1* gene with other Functional Genes predicted by GeneMania

**Table 3.**
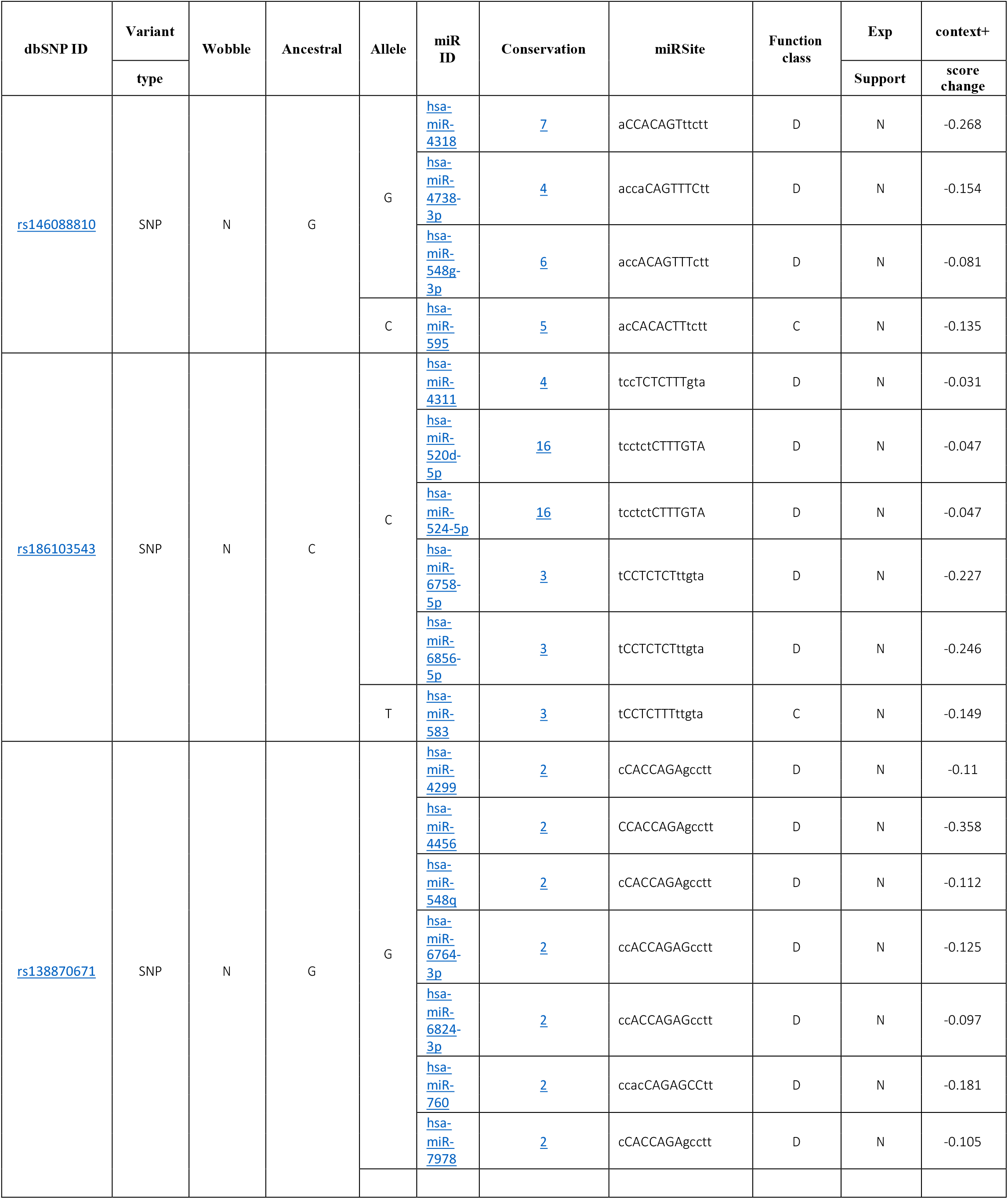
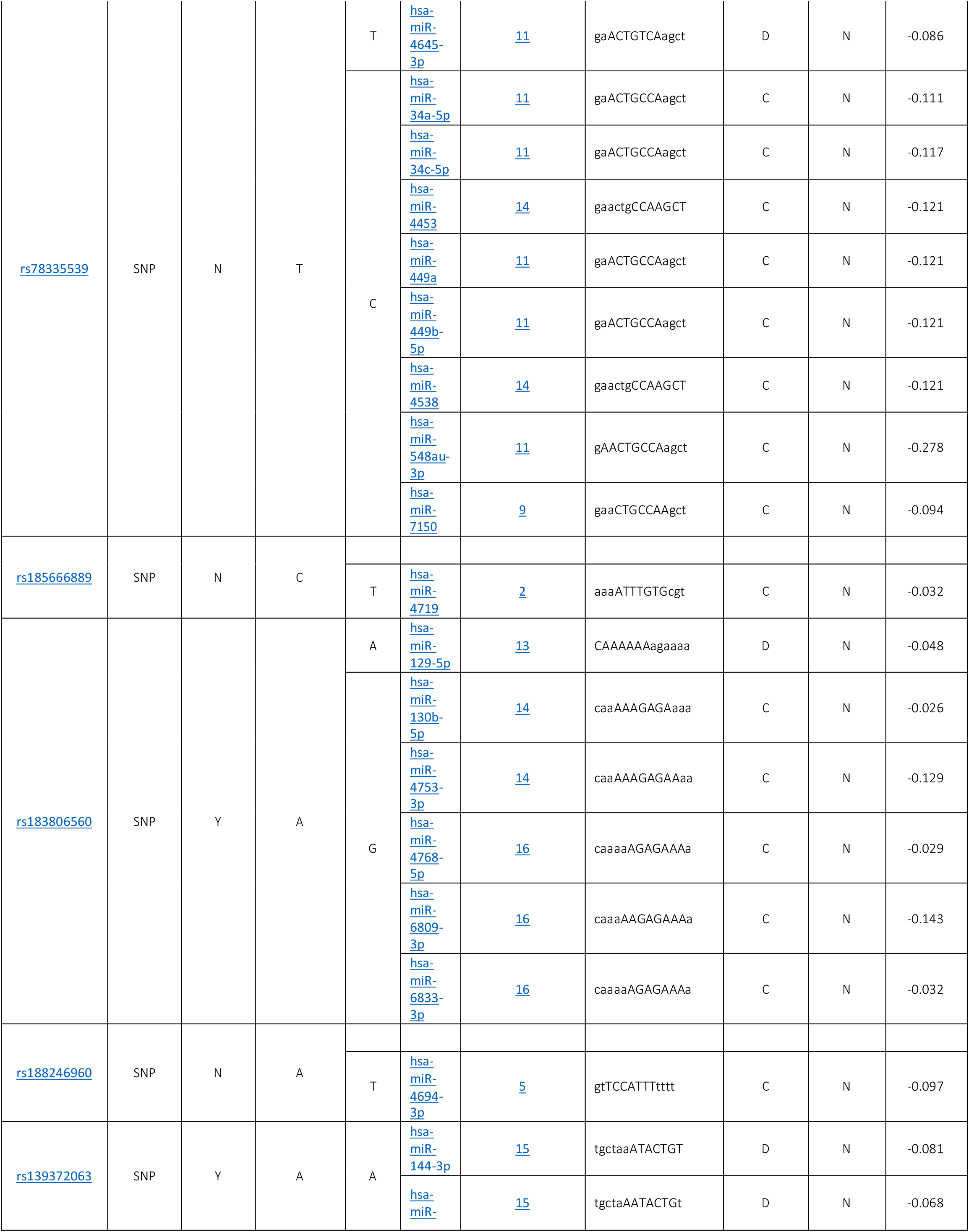
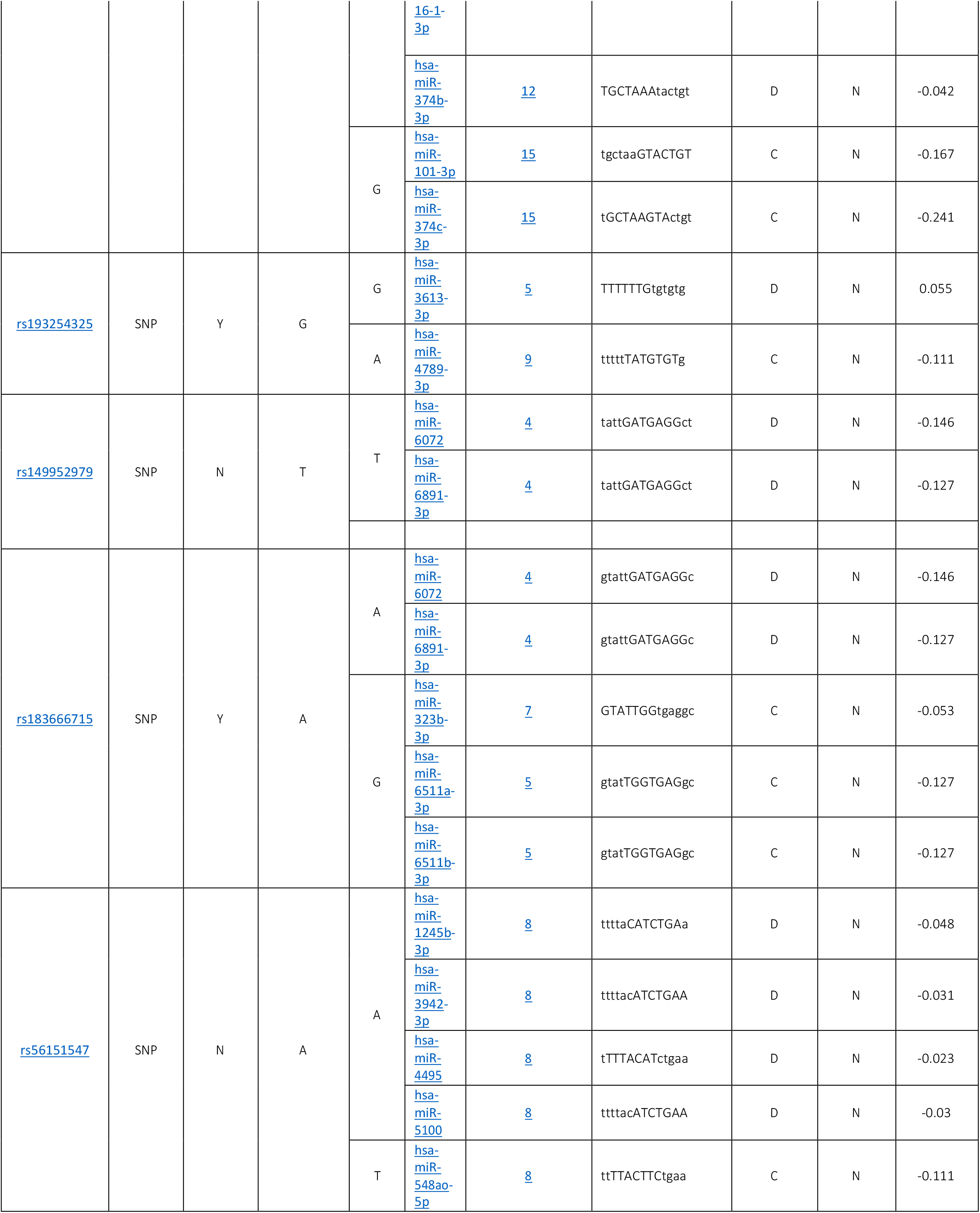
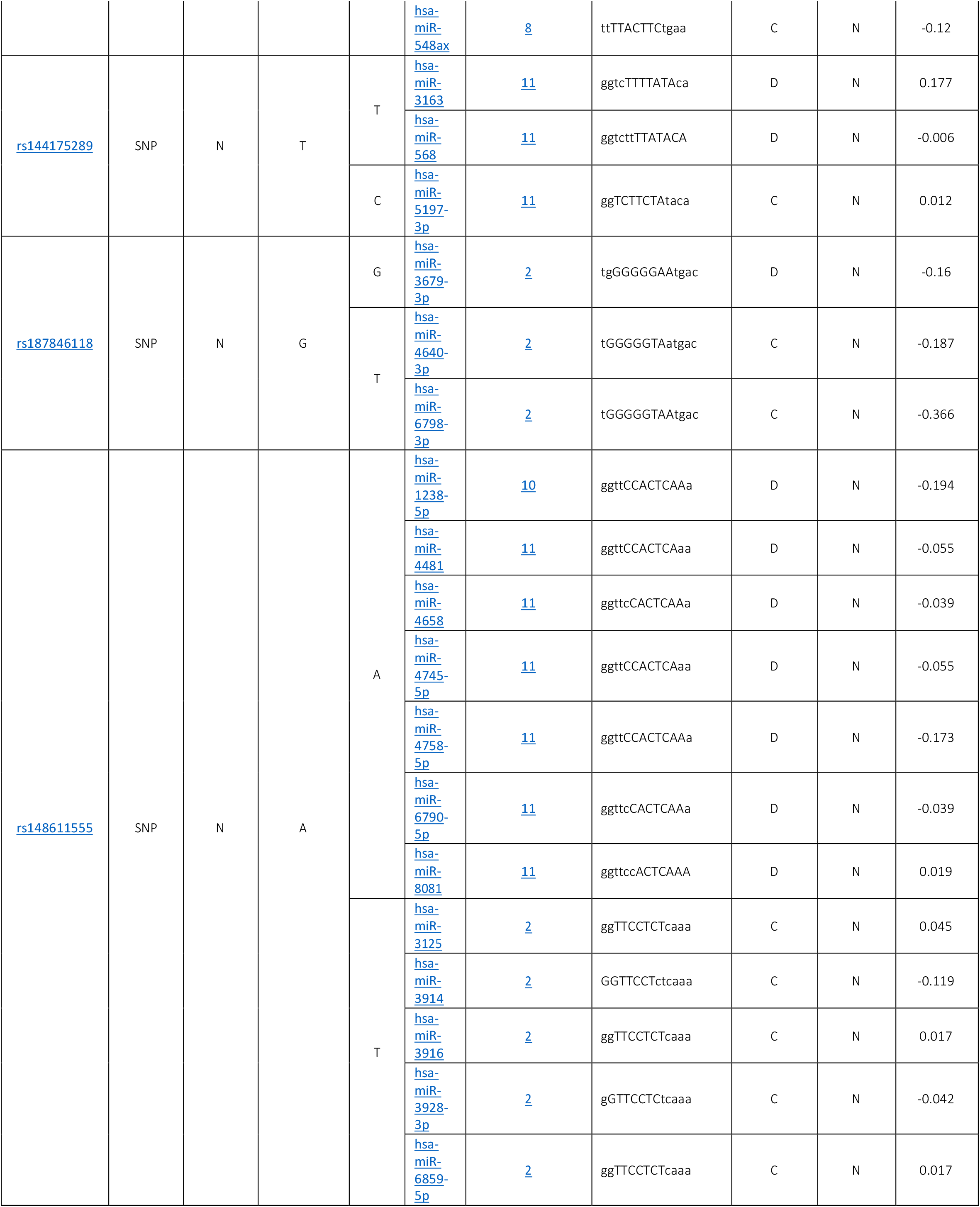
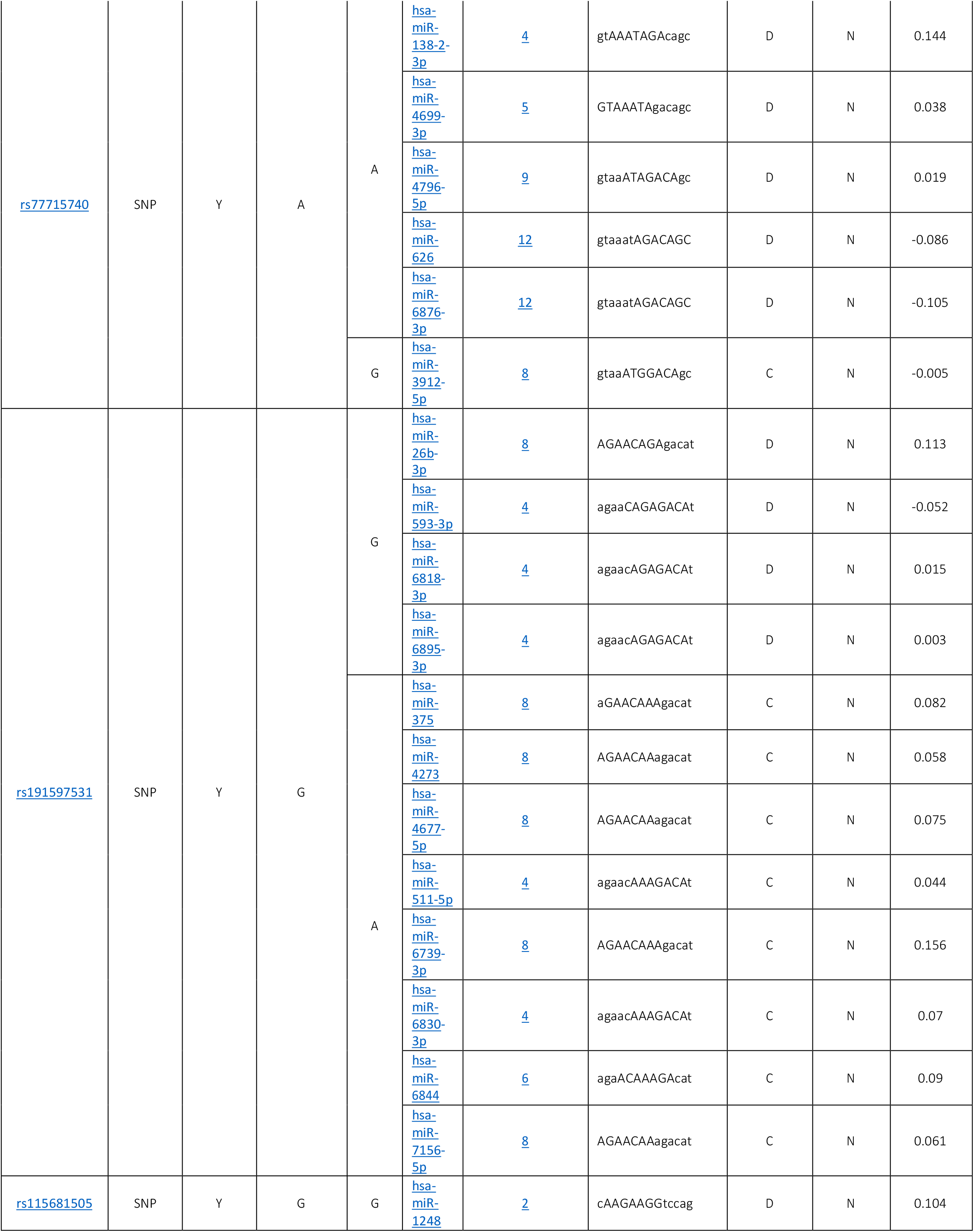
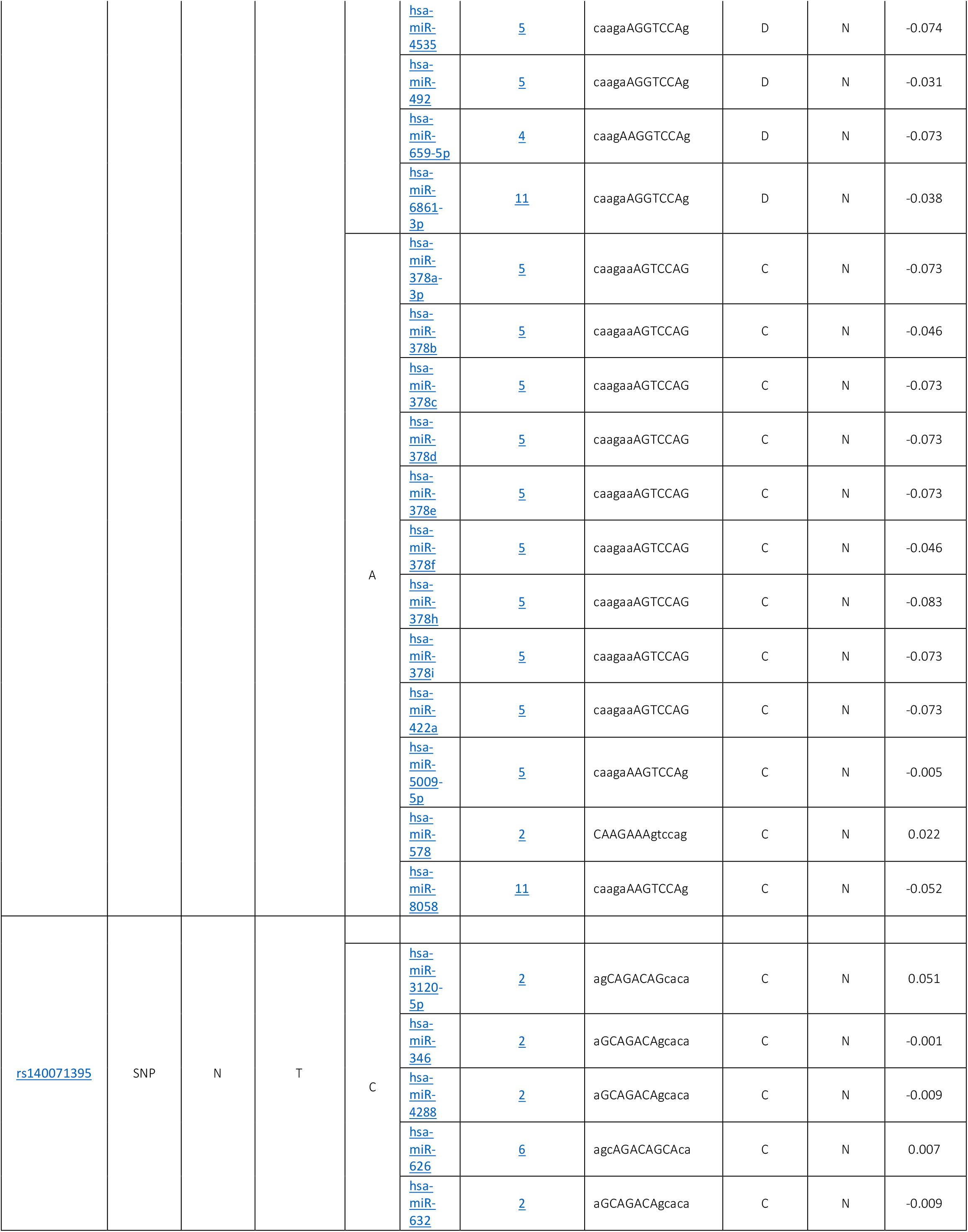
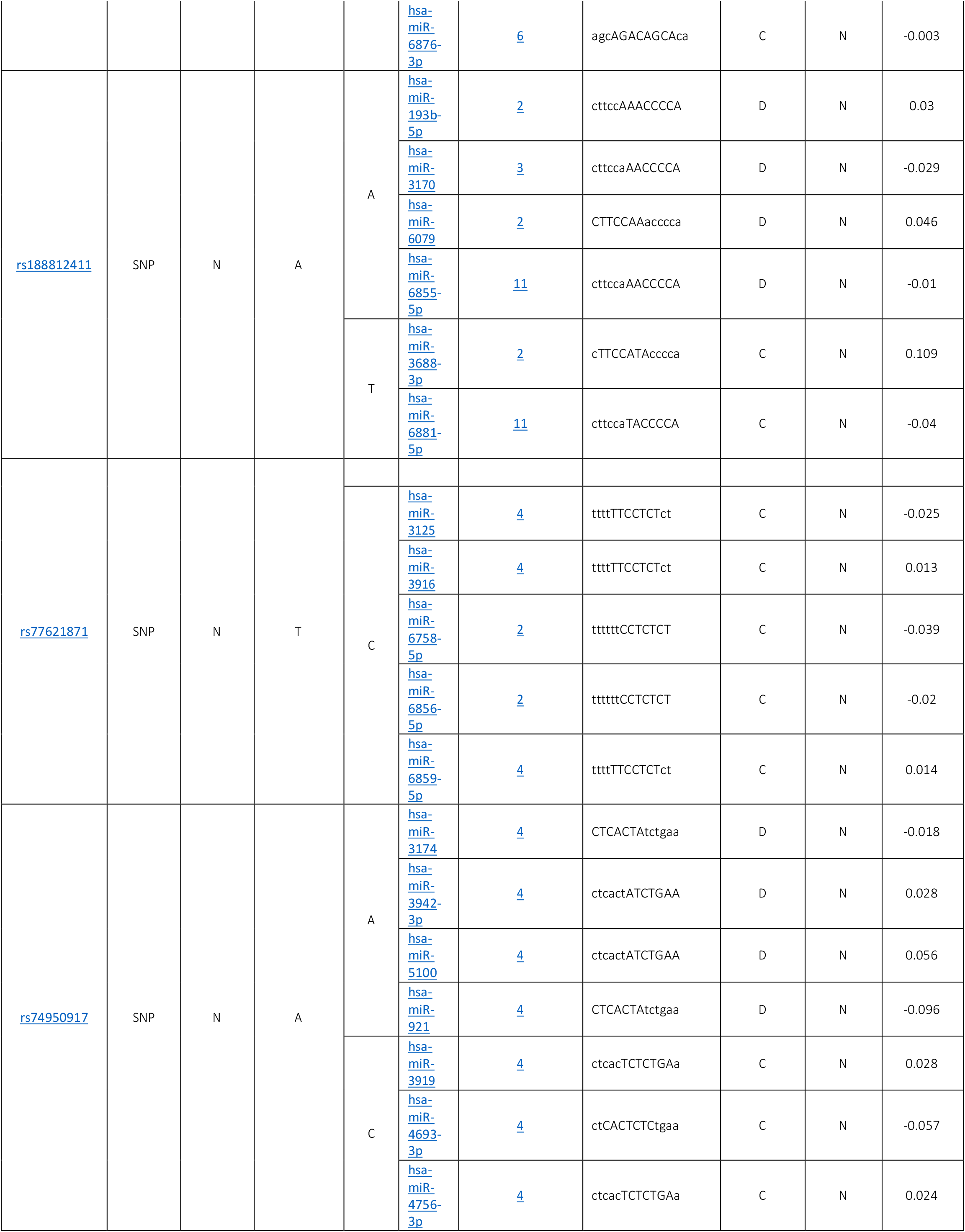
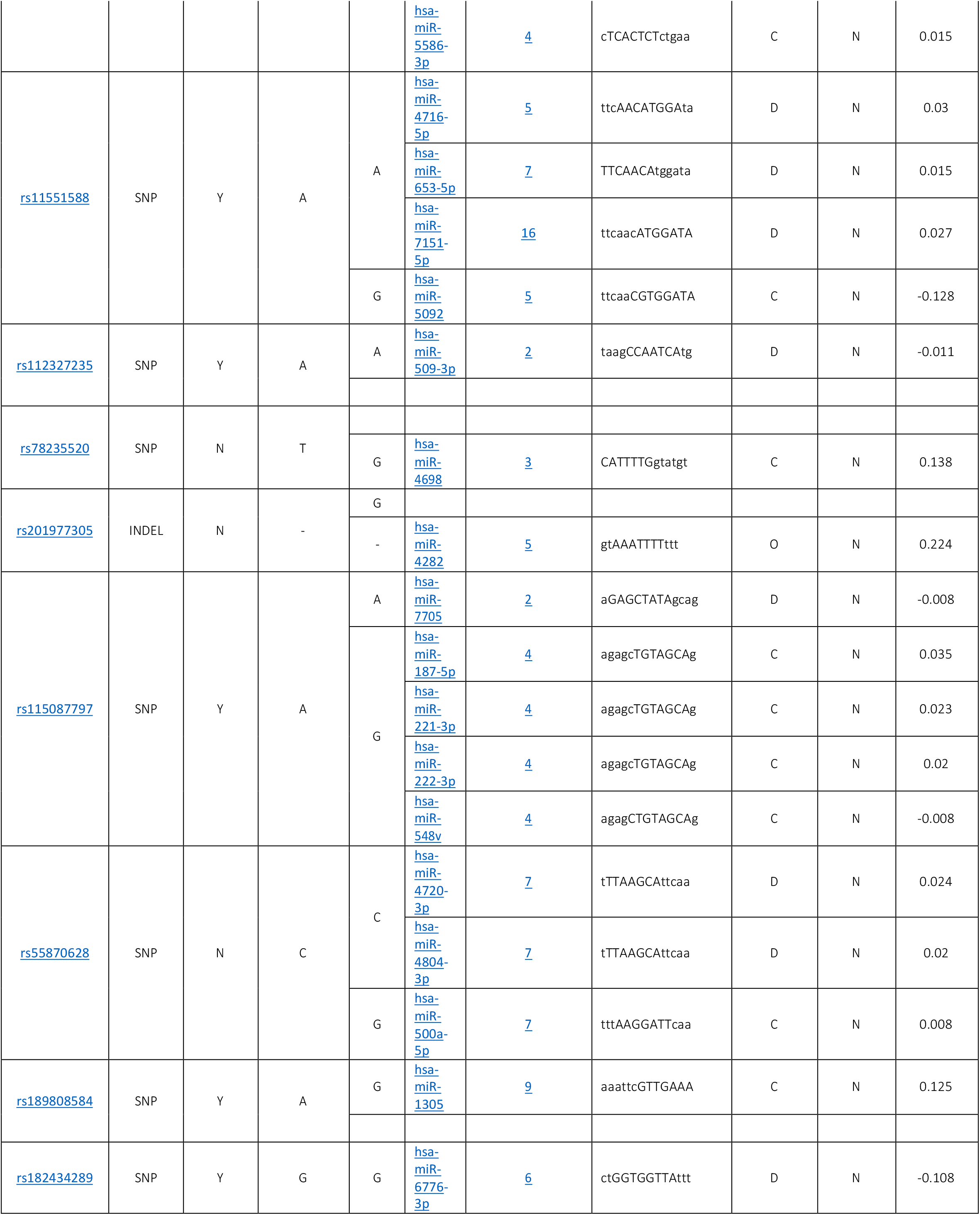
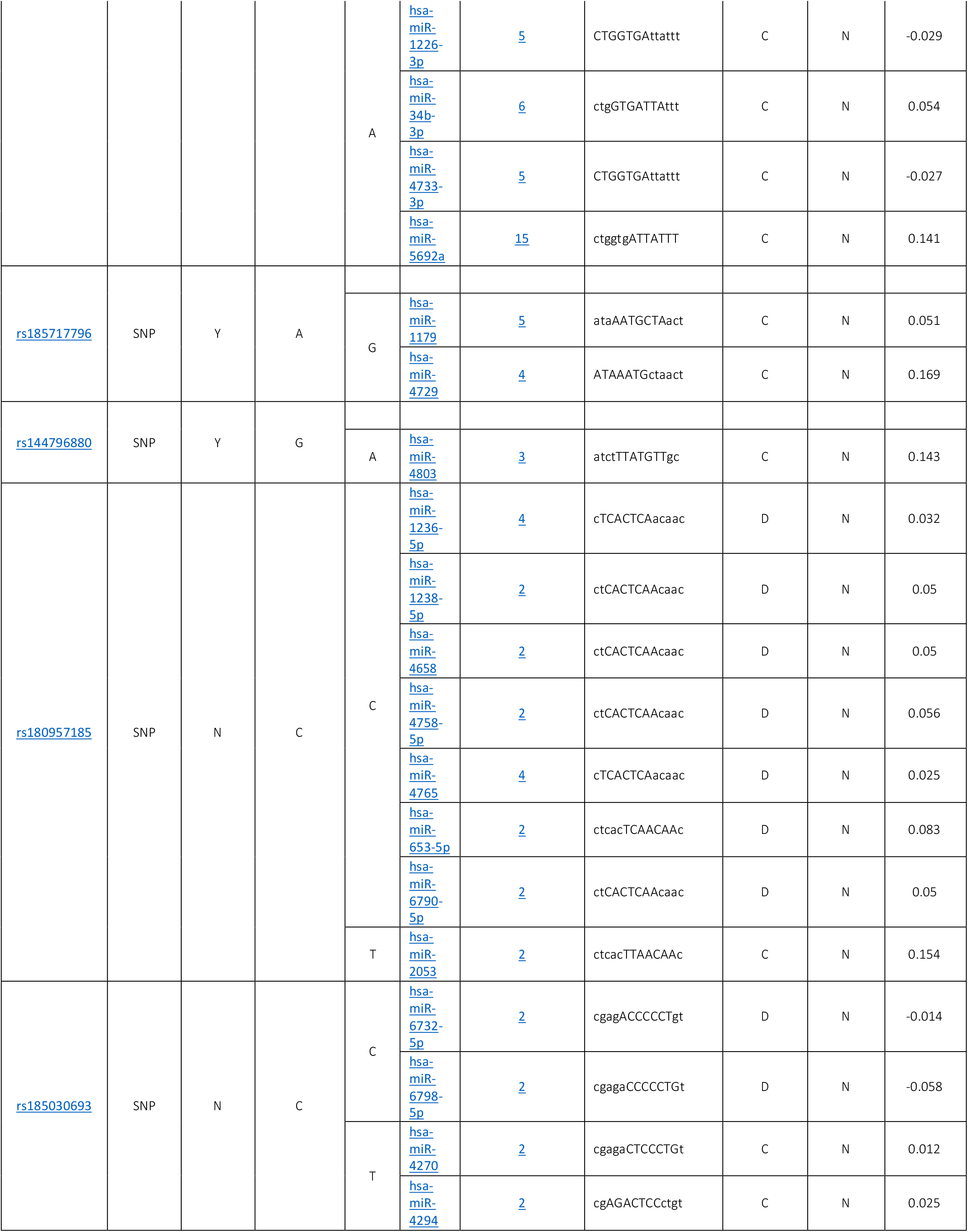
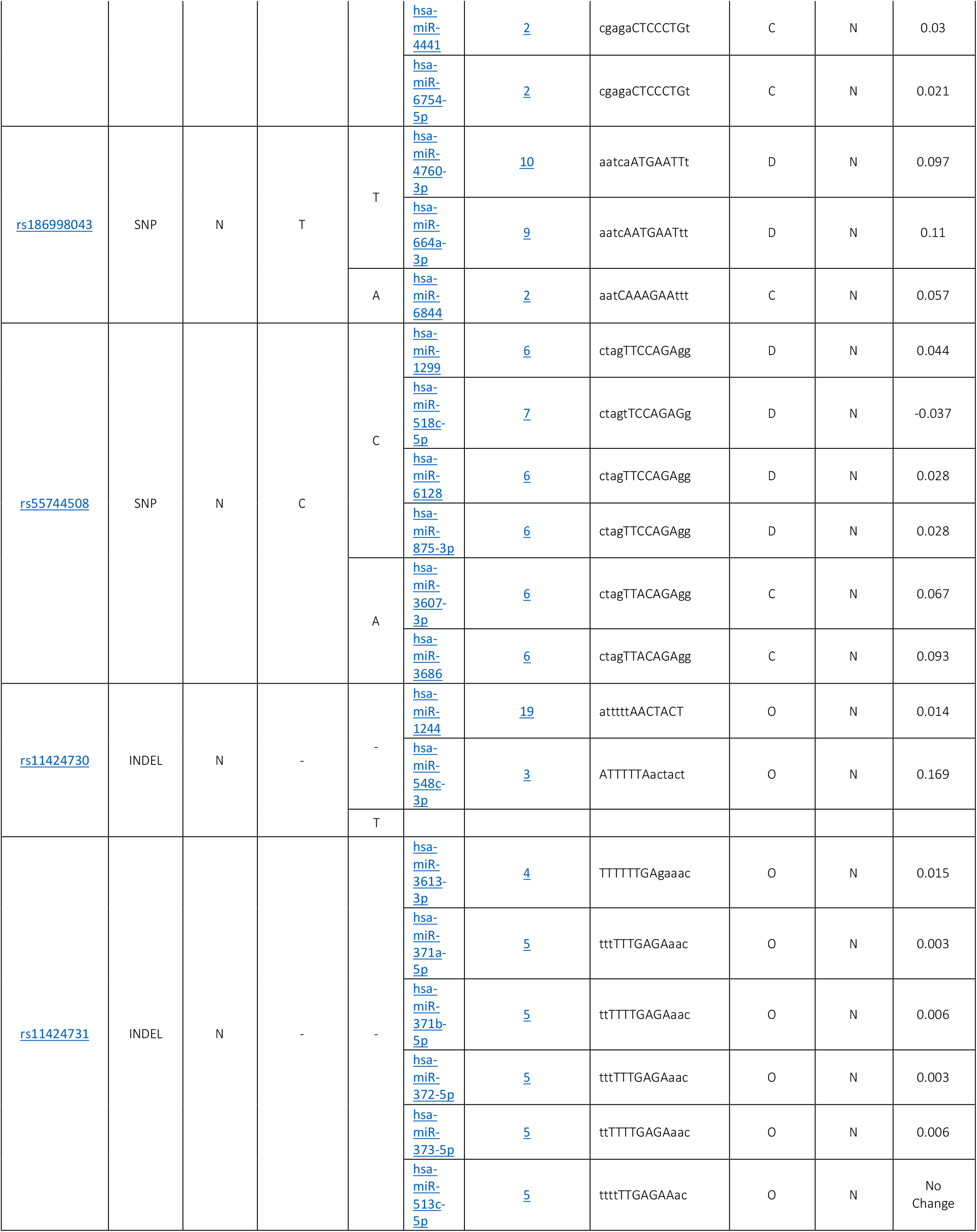
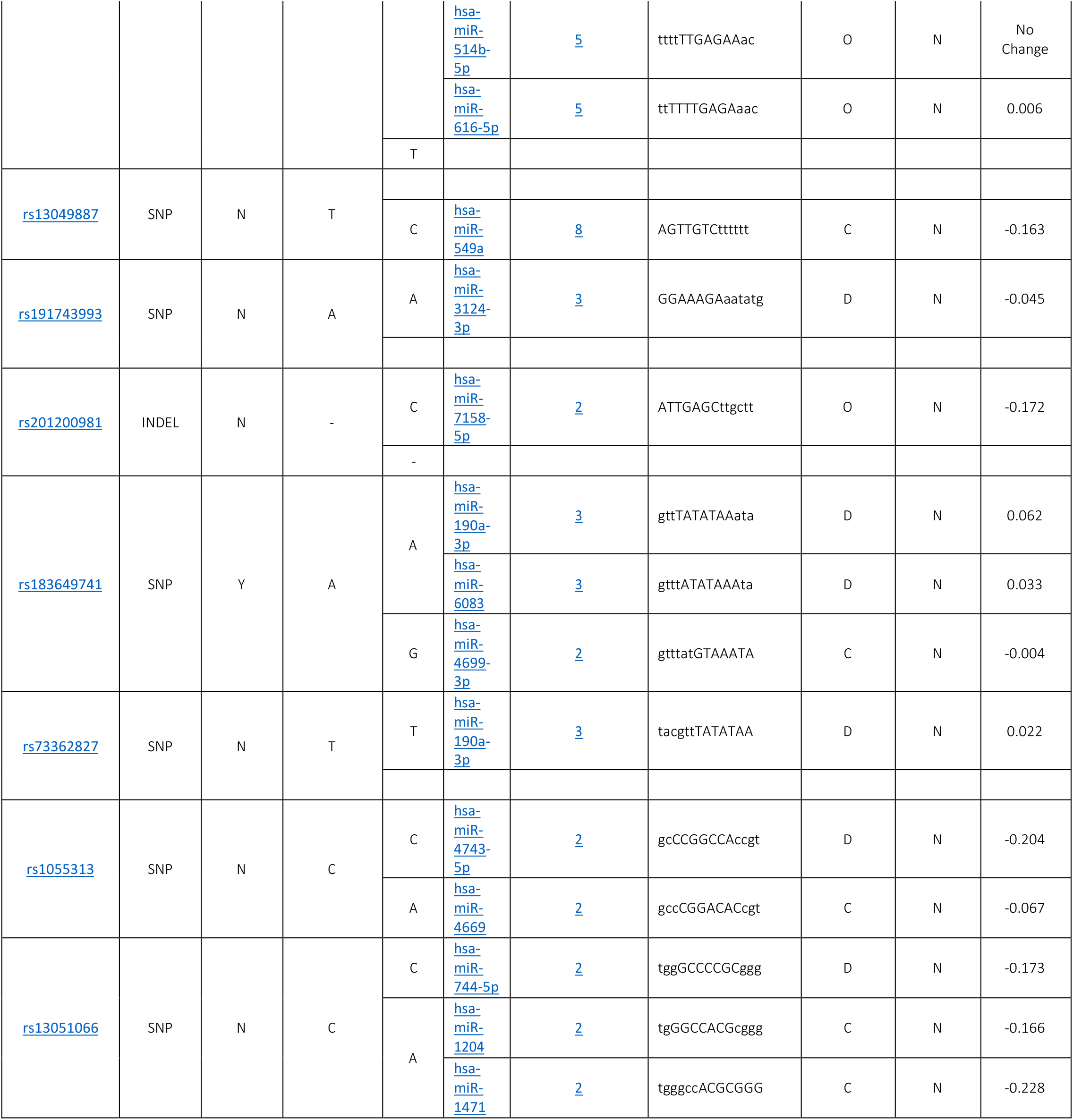
Prediction of SNPs at the 3’UTR Region of *RUNX1* gene using PolymiRTS database

## DISCUSSION

This study discovered seven damaging SNPs, six of them found to be novel SNPs in *RUNX1* gene while only one SNP found in previous studies. These damaging missense mutations in coding region are detected by using different bioinformatics algorithms and softwares.

G165S is the only SNP mentioned in prior studies in patient with myeloid leukaemia (24). According to our analysis the wild-type residue is very conserved but unlike other mutations it has bigger mutant residue than wild-type residue. The residue is located on the surface of the protein, mutation of this residue can disturb interactions with other molecules or other parts of the protein. The torsion angles for this residue are unusual. only glycine is flexible enough to make these torsion angles, mutation into another residue will force the local backbone into an incorrect conformation and will disturb the local structure. The mutation introduces an amino acid with different properties could affect the local structure and as a consequence affect this binding site and Interaction with DNA and abolish its function.

The novel H459P, Y285C L144P, V155A, N109S and N109H mutations have a very conserved wild-type residues which is differ from mutant residues in size that can cause a possible loss of external interactions which might affect the function of the protein. On other hand, H459P and Y285C mutations introduce a more hydrophobic residues at these position that can result in loss of hydrogen bonds and/or disturb correct folding.

Regarding 3’UTR, out of the 000000 SNPs there were 00000 SNPs found to have an effect on it. 201 functional classes were predicted among the 00000 SNPs; 90 alleles disrupted a conserved miRNA site and 99 derived alleles creating a new site of miRNA and this might result in deregulation of the gene function.

Genemania predicts that *RUNX1* functions as: alpha-beta T cell activation, alpha-beta T cell differentiation, antigen receptor-mediated signaling pathway, B cell receptor signaling pathway, bi-cellular tight junction assembly, cartilage development, CD4-positive, alpha-beta T cell activation, CD4-positive, alpha-beta T cell differentiation, CD8-positive, alpha-beta T cell activation, cellular response to steroid hormones stimulus, connective tissue development, hormone-mediated signaling pathway, interleukin-2 production, intracellular estrogen receptor signaling pathway, intracellular receptor signaling pathway, intracellular steroid hormones receptor signaling pathway, leukocyte cell-cell adhesion, lymphocyte differentiation, myeloid cell differentiation, negative regulation of alpha-beta T cell activation, negative regulation of alpha-beta T cell differentiation, negative regulation of CD4-positive, alpha-beta T cell activation, negative regulation of cell activation, negative regulation of cell adhesion, negative regulation of cell-cell adhesion, negative regulation of hemopoiesis, negative regulation of immune system process, negative regulation of leukocyte activation, negative regulation of leukocyte cell-cell adhesion, negative regulation of leukocyte differentiation, negative regulation of lymphocyte activation, negative regulation of lymphocyte differentiation, negative regulation of T cell activation, negative regulation of T cell differentiation, peripheral nervous system development, positive regulation of alpha-beta T cell activation, positive regulation of alpha-beta T cell differentiation, positive regulation of cell activation, positive regulation of cell adhesion, positive regulation of cell-cell adhesion, positive regulation of hemopoiesis, positive regulation of leukocyte activation, positive regulation of leukocyte cell-cell adhesion, positive regulation of leukocyte differentiation, positive regulation of lymphocyte activation, positive regulation of lymphocyte differentiation, positive regulation of T cell activation, positive regulation of T cell differentiation, positive regulation of vasculature development, regulation of alpha-beta T cell activation, regulation of alpha-beta T cell differentiation, regulation of antigen receptor-mediated signaling pathway, regulation of CD4-positive, alpha-beta T cell activation, regulation of CD4-positive, alpha-beta T cell differentiation, regulation of CD8-positive, alpha-beta T cell activation, regulation of hemopoiesis, regulation of interleukin-2 production, regulation of intracellular steroid hormone receptor signaling pathway, regulation of leukocyte cell-cell adhesion, regulation of leukocyte differentiation, regulation of lymphocyte differentiation, regulation of myeloid cell differentiation, regulation of T cell activation, regulation of T cell differentiation, regulatory T cell differentiation, RNA polymerase II transcription regulator complex, steroid hormones mediated signaling pathway, T cell differentiation and transcription regulator complex.

## CONCLUSION

This study revealed 7 damaging SNPs, 6 novel nsSNP out of them in the *RUNX1* gene that leads to AML, by using different bioinformatics tools. Also, 23 functional classes were predicted in 8 SNPs in the 3’UTR, among them, 6 alleles disrupted a conserved miRNA site and 16 derived alleles created a new site of miRNA. This might result in the de regulation of the gene function. Hopefully, these results will help in genetic studying and diagnosis of AML improvement.

## ACKNOWLEDGMENT

Authors wish to acknowledgment the enthusiastic cooperation of Alneelain Stem Cell Center & Africa City of Technology, Khartoum - Sudan.

## CONFLICT OF INTEREST

Authors declare that there is no conflict of interest regarding publication of this paper.

## DATA AVAILABILITY

All relevant data used to support the findings of this study are included within the manuscript and supplementary information files.

## Notes

### Competing Interest Statement

The authors have declared no competing interest.

